# ERVcancer: a web resource designed for querying activation of human endogenous retroviruses across major cancer types

**DOI:** 10.1101/2024.09.02.610762

**Authors:** Xiaoyun Lei, Song Mao, Yinshuang Li, Shi Huang, Jinchen Li, Wei Du, Chunmei Kuang, Kai Yuan

**Author notes:** These authors contributed equally (Xiaoyun Lei, Song Mao). Correspondence (K.Y.).

## Abstract

Human endogenous retroviruses (HERVs) compose approximately 8% of the human genome, co-opted into the dynamic regulatory network of cellular potency in early embryonic development. In recent studies, resurgent HERVs’ transcriptional activity has been frequently observed in many types of human cancers, suggesting their potential functions in the occurrence and progression of malignancy. However, a web resource dedicated to querying the relationship between activation of HERVs and cancer development is lacking. Here, we have constructed a database to explore the sequence information, expression profiles, survival prognosis, and genetic interactions of HERVs in diverse cancer types. Our database currently incorporates RNA sequencing (RNA-seq) data of 580 HERVs across 16246 samples, comprising 151 early embryonic data from the Gene Expression Omnibus (GEO) database, 8051 human adult tissues’ data from the Genotype-Tissue Expression (GTEx) project, 932 cancer cell lines’ data from the Cancer Cell Line Encyclopedia (CCLE) project, 6478 tumoral and 634 normal tissue samples’ data from The Cancer Genome Atlas (TCGA) project. The primary goal is to provide an easily accessible and user-friendly database for professionals in the fields of bioinformatics, pathology, pharmacology, and related areas, enabling them to efficiently screen the activity of HERVs of interest in normal and cancerous tissues and evaluate the clinical relevance. The ERVcancer database is available at http://kyuanlab.com/ervcancer/.

## Introduction

Human endogenous retroviruses (HERVs) are remnants of ancient retroviral infections integrated into the human germline cells over millions of years (de Parseval and Heidmann, 2005; Jern and Coffin, 2008). These retroviral sequences have been transmitted and fixed across generations, constituting approximately 8% of the human genome, a portion that is even larger than that of protein-coding sequences (Lander et al., 2001; Nurk et al., 2022). Most of HERVs reside within heterochromatin in somatic cells, where they are predominantly silenced by diverse epigenetic mechanisms, including DNA methylation, H3K9me3 and H3K27me3 histone modifications (Rowe et al., 2010; Groh and Schotta, 2017; Cosby et al., 2019; Zhao et al., 2023). However, during the profound epigenetic resetting in early embryogenesis (Zheng et al., 2016; Du et al., 2022), or in tumorigenesis with altered epigenetic landscapes (Berdasco and Esteller, 2010; Saghafinia et al., 2018; Michalak et al., 2019), many endogenous retroviruses often get activated (Göke et al., 2015; Grow et al., 2015; Kong et al., 2019; He et al., 2021; Dopkins and Nixon, 2024; Liang et al., 2024), impacting the transcription of host genes or even producing retroviral products to modify the cellular regulatory processes (Wang et al., 2014; Zhang et al., 2019; Ai et al., 2022; Fueyo et al., 2022; Yu et al., 2022; Sakashita et al., 2023; Gong et al., 2024).

The activation of specific endogenous retroviruses (ERVs) has been functionally linked to various biological processes, including embryonic development (Grow et al., 2015; Zhang et al., 2019; Asimi et al., 2022; Sakashita et al., 2023), germline specification (Zhou et al., 2022), neuronal differentiation (Johansson et al., 2020; Padmanabhan Nair et al., 2021; Yan et al., 2022), innate immunity (Chuong et al., 2016; Li et al., 2023; Russ and Iordanskiy, 2023), and aging (Liu et al., 2023; Wang et al., 2024). For instance, human endogenous retrovirus H (HERVH) is dynamically transcribed during the early stages of human embryonic development, promoting the characteristics of a naive-like state of human embryonic stem cells (hESCs) (Wang et al., 2014; Ai et al., 2022). More recently, abnormal activation of HERVs has been detected in many types of cancer, such as cervical (Soleimani-Jelodar et al., 2024), gallbladder (Wang et al., 2022), pancreatic (Cortesi et al., 2024), colorectal (Zhang, D.Z. et al., 2024), hepatocellular (Zhou et al., 2021), urothelial cell (Yu et al., 2014), breast (Wang-Johanning et al., 2014), and testicular cancers (Gimenez et al., 2010; Li et al., 2024). Specifically, the activation of human endogenous retrovirus K (HERVK) plays a role in establishing a stem cell niche in glioblastoma (Shah et al., 2023). Our previous study has demonstrated that HERVH transcripts steer the BRD4 coactivation module to promote oncogenicity, and downregulation of HERVH transcript levels effectively inhibits the proliferation of colorectal cells as well as patient-derived colorectal cancer organoids (Li et al., 2022; Yu et al., 2022). Besides, HERVH-derived lncRNA UCA1 plays a crucial role in gastric cancer cell proliferation (Bian et al., 2016; Wang et al., 2019). Of note, activated HERVs can also function as promoters, further driving the expression of numerous oncogenes through a process known as onco-exaptation (Babaian and Mager, 2016; Jang et al., 2019; Attig et al., 2023). Beyond human cancers, abnormal activation of HERVs has also been observed in other diseases, including multiple sclerosis (Dolei, 2018), schizophrenia (Tamouza et al., 2021; Wu et al., 2023a; Wu et al., 2023b; Xue et al., 2023; Yao et al., 2023; Zhang, D.Y. et al., 2024), and type 1 diabetes (Levet et al., 2019). The frequent associations of resurgent HERVs’ transcriptional activity with diverse human diseases highlight their potentials as biomarkers or even future therapeutic targets in clinic.

To date, most of the HERVs-related databases have primarily been used to provide annotations for these repetitive elements and identify conserved sequences, such examples include RepeatMasker (Smit et al., 2013-2015), RepBase (Bao et al., 2015), Dfam (Hubley et al., 2016), ERVd (Paces et al., 2004), and ERVmap (Tokuyama et al., 2018). More recently, Erik Stricker et al. utilized the PDF data extractor (PDE) R package to extract HERV expression information from PubMed literature tables, and established CancerHERVdb (Stricker et al., 2023), a database that allows for the retrieval of relevant literature regarding the expression of specific HERVs in particular cancers. Currently, there are no ready-to-use databases available for easily exploring the expression profiles of different HERVs across multiple cancer tissues or cell lines.

To meet this need, we present ERVcancer, a web resource designed for analyzing transcription of HERVs of interest in the occurrence and progression of human malignancies in different tissues. It not only covers the expression profiles of HERVs at different embryonic developmental stages and in different types of healthy tissues, but also offers differential expression analyses between peritumoral and tumoral samples of various cancer types. Moreover, the web resource allows users to link HERVs’ expression profiles to the clinical parameters of a given cancer type, and to query the potential correlations between the expressions of HERVs and coding genes, providing useful guidance for subsequent functional interrogations of HERVs in cancer progression. Due to the repetitive nature of HERVs, it is challenging to precisely allocate every sequencing reads of HERVs to a unique genomic locus. Therefore, in the current version of the database we only include analyses of bulk expression of each HERV subfamily. ERVcancer is freely available at http://kyuanlab.com/ervcancer/.

## Results

### Data collection and analysis

To feed the ERVcancer database, we collected a substantial amount of RNA-seq data generated using embryonic cells, normal somatic tissues, cancer cell lines, as well as tumoral and matched normal tissues. Specifically, 151 FASTQ-formatted RNA-seq data of human early embryos and embryonic stem cells were gathered from Gene Expression Omnibus (GEO) database(Xue et al., 2013; Yan et al., 2013) (Figure 1A); the expression profiles of HERVs in 8051 samples from 49 different adult tissues were obtained from Genotype-Tissue Expression (GTEx) project (Bogu et al., 2019) (Figure 1B); FASTQ-formatted RNA-seq data of 932 cancer cell lines from Cancer Cell Line Encyclopedia (CCLE) project were downloaded from EMBL-EBI (European Molecular Biology Laboratory-European Bioinformatics Institute) (https://www.ebi.ac.uk/) (PRJNA523380) (Ghandi et al., 2019) (Figure 1C); and 7193 BAM-formatted read alignment files (for GRCh38/hg38) of 12 different cancer types in The Cancer Genome Atlas (TCGA) were accessed through dbGaP (database of Genotypes and Phenotypes) (https://www.ncbi.nlm.nih.gov/gap/) accession number phs00178.v11.p8 and downloaded from the Genomic Data Commons (GDC) data portal site (https://portal.gdc.cancer.gov/) using the GDC Data Transfer Tool (Figure 1D). Among these BAM-formatted files, 634 normal and 6478 primary tumor samples were kept, and Formalin Fixed Paraffin-Embedded (FFPE) samples were excluded to avoid potential complications of RNA degradation.

**Figure 1.**
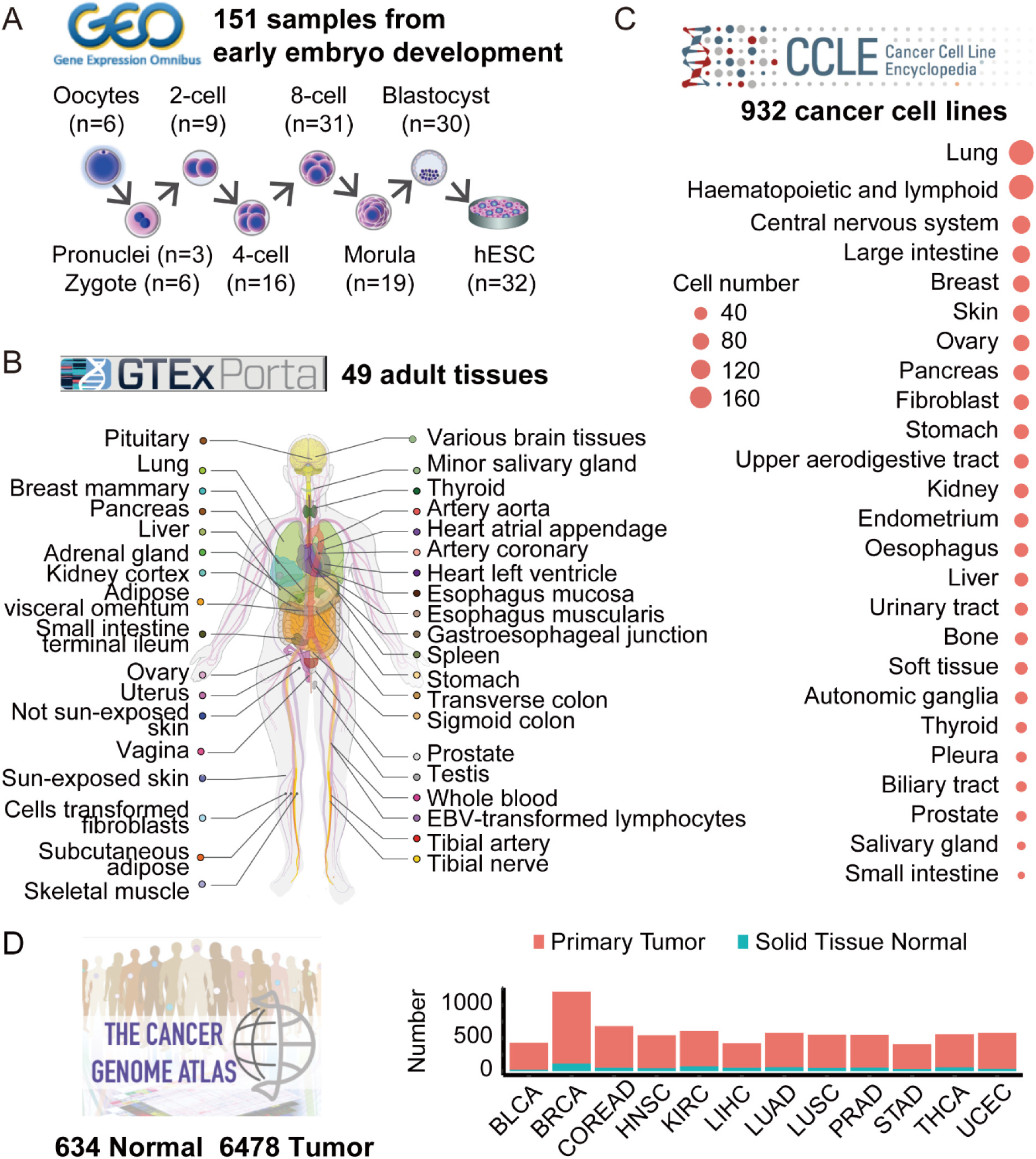
RNA-seq datasets included in ERVcancer. (A) RNA-seq datasets from different stages of human embryos obtained from GEO (accession number GSE44183, GSE36552). (B) RNA-seq datasets from various tissue sites obtained from GTEx. (C) RNA-seq datasets of cancer cell lines obtained from CCLE. The dot size represents the number of cell lines for each tissue type. (D) RNA-seq datasets of various cancer types obtained from TCGA. The height of bars represents the number of samples for each cancer type that were analyzed.

To quantify the expression of HERVs, we first aligned FASTQ-formatted RNA-seq reads to the reference genome (for GRCh38/hg38) using STAR (Dobin et al., 2013), then counted the aligned reads using featureCounts (Liao et al., 2014). Due to HERVs are present in many hundreds to millions of copies within the genome, multiple mapped reads are kept to avoid missing HERVs’ expression information. While assigning these reads to the best alignment location is a straightforward method to resolve HERV-derived reads, it may not always provide accurate results for each individual copy (Treangen and Salzberg, 2012; Goerner-Potvin and Bourque, 2018). To solve this problem, reads mapping to any HERV copy in the genome are assigned to a unique HERV subfamily (He et al., 2021). This strategy is instrumental in reducing the error rate in multimapping read allocation. To verify the robustness of our pipeline for HERVs quantification, we compared the results with that generated by the REdiscoverTE, which is a Salmon-based tool for quantifying transposable elements using a slightly different algorithm for multiple mapping reads allocation (Patro et al., 2017; Kong et al., 2019; Deschamps-Francoeur et al., 2020). Our pipeline exhibited high consistency with REdiscoverTE for transposable element quantification (Figure S1A), and the subsequent statistical analyses based on the quantifications generated by these two methods yielded similar results (Figure S1B-S1E, 3C-3F).

To examine whether HERVs exhibit transcriptional activation in a given cancer type, we employed a methodology similar to differential expression genes analysis between tumor and normal samples (Love et al., 2014), and identified HERV subfamilies that show aberrant expression in tumors. We included 12 tumor types with a considerable number of normal tissue samples from the TCGA in the current version of ERVcancer database. Due to a large computational workload involved in this step, we pre-ran the differential analysis for each cancer type to prevent redundant calculations, so that users can directly access these results from the web resource.

To investigate whether the expression levels of HERVs affect the prognosis of cancer patients, we collected the clinical data for the TCGA samples (Liu et al., 2018), including tumor stage, histological type, and survival outcome, and incorporated into the database. For a specific HERV, users can choose median, quartile, or customized threshold to divide tumor cases into groups based on its standardized expression (CPM, counts per million). Subsequently, the database server will perform calculations and return the results, with the information on whether there is a difference in prognosis between patients’ groups stratified by the expression level of the selected HERV.

To data-mine the potential regulatory relationships between HERVs and the coding genes, we computed the correlation coefficients based on their expressions. First, genes’ normalized counts of TCGA patient tumor tissue samples were collected via the UCSC Xena platform (https://xena.ucsc.edu) (Goldman et al., 2020). Second, the standardized expressions of HERVs and the coding genes were merged. Third, pairwise correlations between each HERVs and coding genes were calculated. We then included the correlation coefficients in the ERVcancer database, so that users can identify potential regulations according to the strength of the correlation.

### Web interface and graphic visualization

Raw datasets were organized into tables using SQLite3. Users can freely access expression profiles and the extended analysis results through the web interface, and download the results in the form of tables and images to their local devices (Figure 2A).

**Figure 2.**
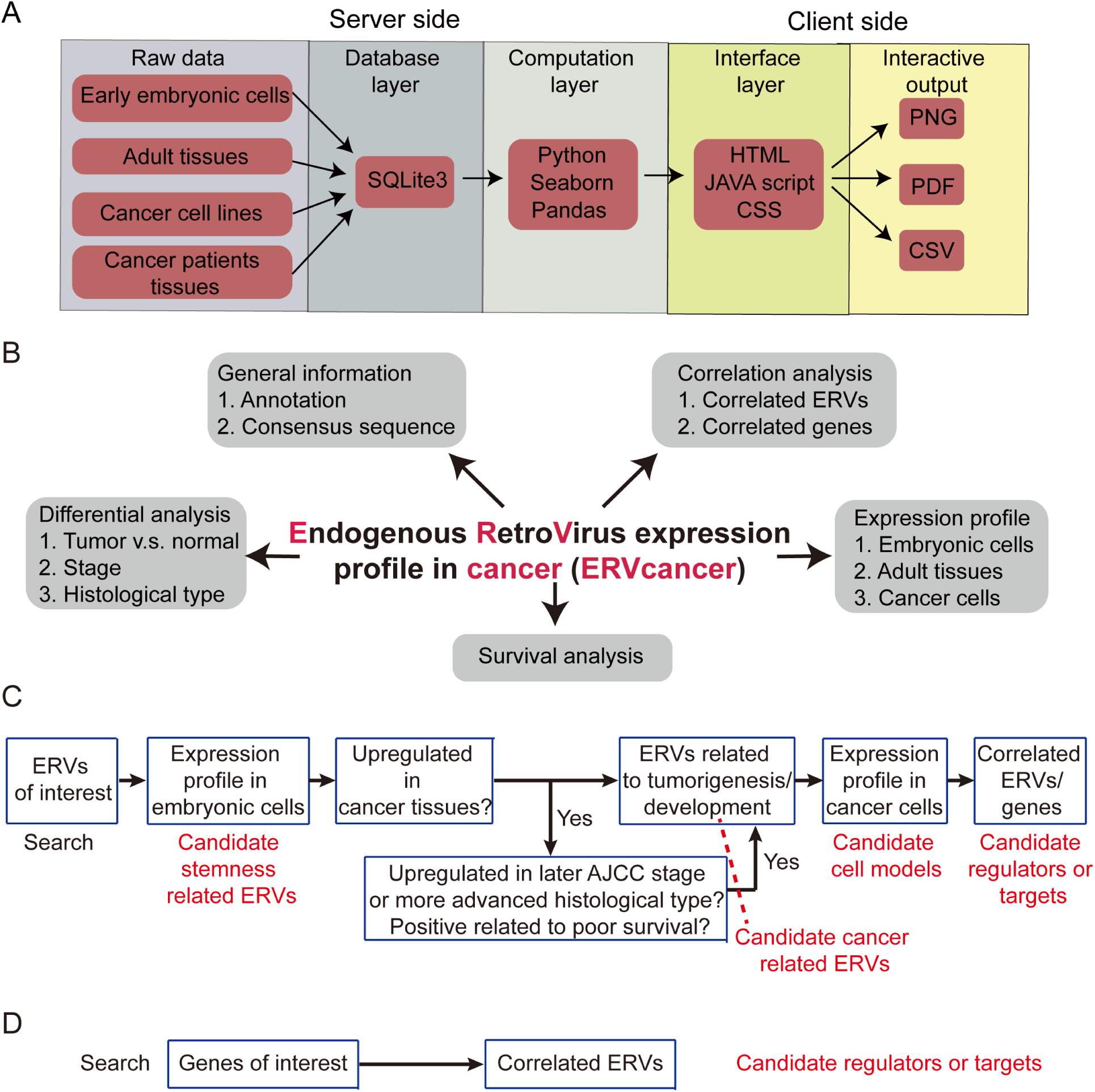
Overview of ERVcancer. (A) Schema describing data processing and display for the ERVcancer visualization tool. (B) Functional modules of ERVcancer, including General information, Expression profile, Differential analysis, Survival analysis, and Correlation analysis. (C-D) The general workflow for querying a HERV (C) or a gene (D) of interest.

The features of the ERVcancer database are categorized into five main tabs: General information, Expression profile, Differential analysis, Survival analysis, and Correlation analysis (Figure 2B). A typical workflow for utilizing ERVcancer functionalities is shown in Figure 2C. For a given HERV of interest, users can first access the General tab to obtain its basic information, such as its classification and consensus DNA sequence. Then, users can investigate its expression profile in embryonic cells to assess its association with pluripotency. Subsequently, users can examine whether it exhibits specific expression patterns in certain cancer type and whether its expression correlates with tumor prognosis. Once a candidate tumor-associated HERVs is identified, its associated genes can be retrieved by correlation analyses to construct a putative regulatory network. On the other hand, users can also search for genes of interest to identify their correlated HERV subfamily in a specific cancer type, obtaining clues about potential regulatory relationships between genes and HERVs (Figure 2D).

### Example of application

#### (HERVH-int as an example)

In this section, we will use “HERVH-int” as a query to provide an intuitive understanding of the performance of our database and its potential applications in the studies of HERVs during tumorigenesis.

Firstly, enter “HERVH-int” in the search bar on the main page. After submitting the query by clicking the GO button, the information listed in the General tab will be returned, displaying annotations and its consensus sequence, which can be used to guide primers’ design in subsequent experiments.

Next, users can click on the Expression profile tab. By selecting sample types from the dropdown options (including early human embryonic cells, human adult tissues, and cancer cell lines), its expression profile across different samples can be retrieved. HERVH-int is highly expressed in hESCs as previously reported (Göke et al., 2015) (Figure 3A), and is abnormally activated in several colorectal cancer cell lines, such as SNU1040, LS123, and HCT116 (Figure 3B). Users can select appropriate cell lines for the follow-up experiments based on the relative expression levels. For example, HERVH-int is highly expressed in HCT116 but not in SW480, so that HCT116 may be suitable for HERVH-int RNAi experiments while SW480 can be selected for CRISPR activation (CRISPRa) assays.

**Figure 3.**
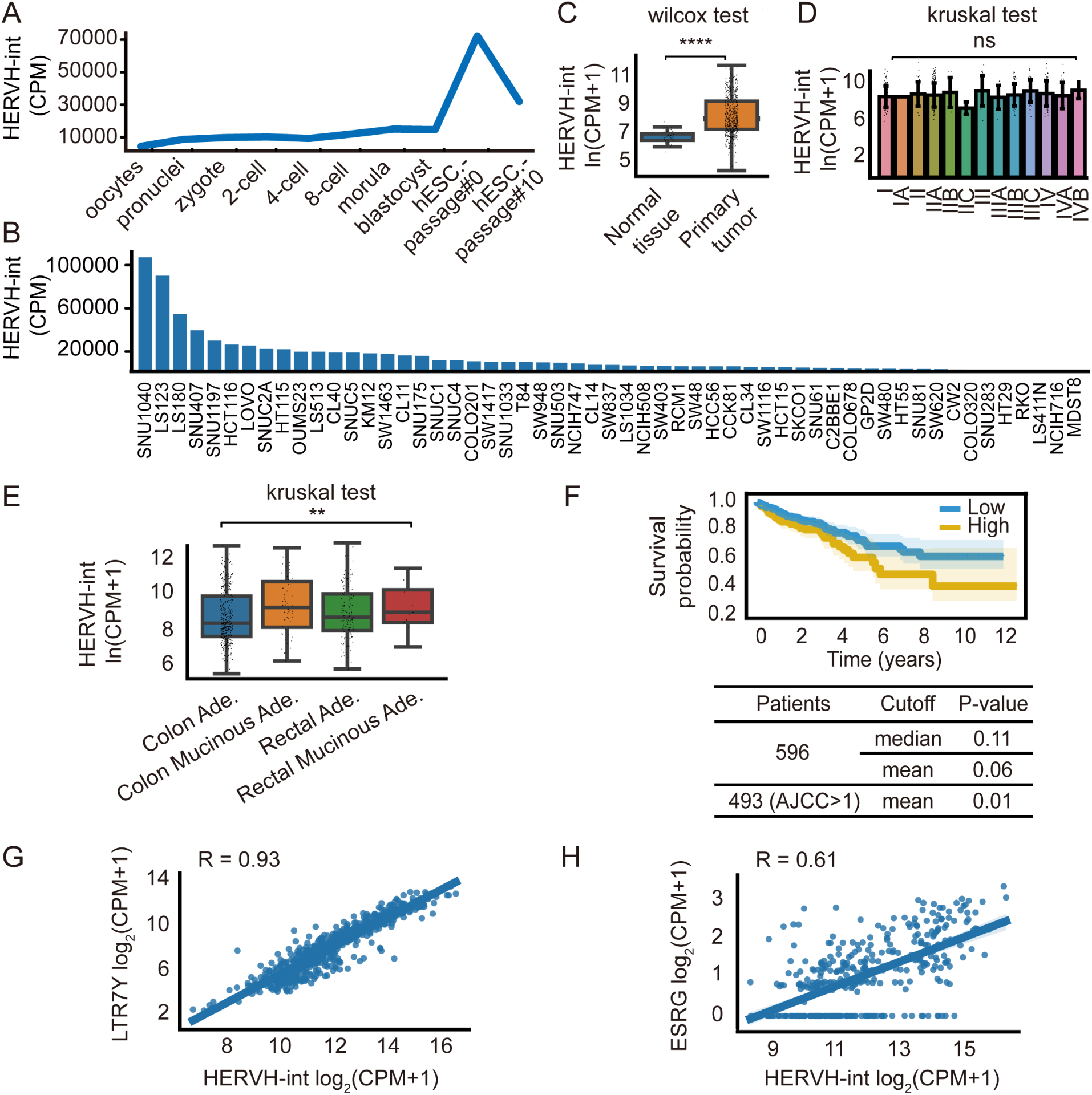
Examples of application using HERVH-int as a query. (A-B) The expression profile of HERVH-int across early embryonic developmental stages (A) and in colorectal cancer cell lines (B). (C-E) Differential expression analysis of HEVH-int among normal and cancer tissues (C), different tumor stages (D), or different histological types (E). Ade. is the abbreviation of adenocarcinoma. (F) Overall survival analysis, grouped by HERVH-int’s expression levels. (G-H) Pairwise correlation analysis of HERVH-int versus other HERVs (G) or genes (H).

Then, users can click on the Differential expression tab. By selecting tumor types and grouping methods (including tumor vs. normal, different stages, and different histological types) from the dropdown options, one can determine if there are any differences in expression levels across different groups. For example, HERVH-int is highly expressed in colorectal cancer samples (Figure 3C), exhibiting statistical differences among different histological types but not tumor stages of colorectal cancers (Figure 3D-3E).

After that, users can click on the Survival analysis tab. By selecting tumor types and different cutoffs (including median, quantile, and custom), one can assess whether the expression of the HERV is associated with the prognosis of a certain type of cancer. In colorectal cancer patients, higher expression of HERVH-int is associated with a poorer prognosis, especially for those with AJCC pathologic tumor stage > 1 (Yu et al., 2022) (Figure 3F).

Finally, users can click on the Correlation tab. By selecting cancer types, one can examine the correlations between the expressions of the queried HERV and coding genes or other HERV elements. The results are sorted in descending order based on the correlation coefficients by default. Genes manifested high correlation may serve as potential upstream or downstream regulatory factors for the queried HERV (Figure 3G-3H).

### Case study

#### (KZFPs and ERVs)

Previous researches have revealed that many Krüppel-associated box domain-containing zinc finger proteins (KZFPs) exhibit the ability to recognize transposable elements (TEs)-embedded sequences as genomic targets, thus serving as TE-controlling repressors (Najafabadi et al., 2015; Helleboid et al., 2019; de Tribolet-Hardy et al., 2023). Yet, a comprehensive analysis of the pairwise relationships between a particular KZFP and a particular HERV subfamily has not been conducted. Based on our ERVcancer database, we performed a pan-cancer survey to investigate the correlations between the expression profiles of KZFPs and HERVs.

Firstly, expression profiles of HERVs and KZFP genes from 12 different cancer types were aggregated. Then, the pan-cancer correlations of the expressions between KZFPs and HERVs were calculated. For each pair of KZFP-HERV, only significantly correlated pairs were retained. Finally, we tabulated the number of KZFPs targeting each HERV subfamily (Supplementary Table 1), and the top 10 HERV subfamilies with the highest counts of positively and negatively correlated KZFPs are shown in Figure 4A. We observed that members of the HERVH subfamily, including LTR7Y and HERVH-int, were negatively correlated with the expression of numerous KZFPs (Figure 4B). We further plotted the top 20 KZFPs that were negatively correlated with LTR7Y or HERVH-int (Figure 4C-4D). Cytoscape illustration and venn diagram showed that there was a significant overlap between KZFPs negatively correlated with LTR7Y or HERVH-int (Figure 4E-4F), indicating that these two elements were similarly regulated by a set of KZFPs during tumorigenesis.

**Figure 4.**
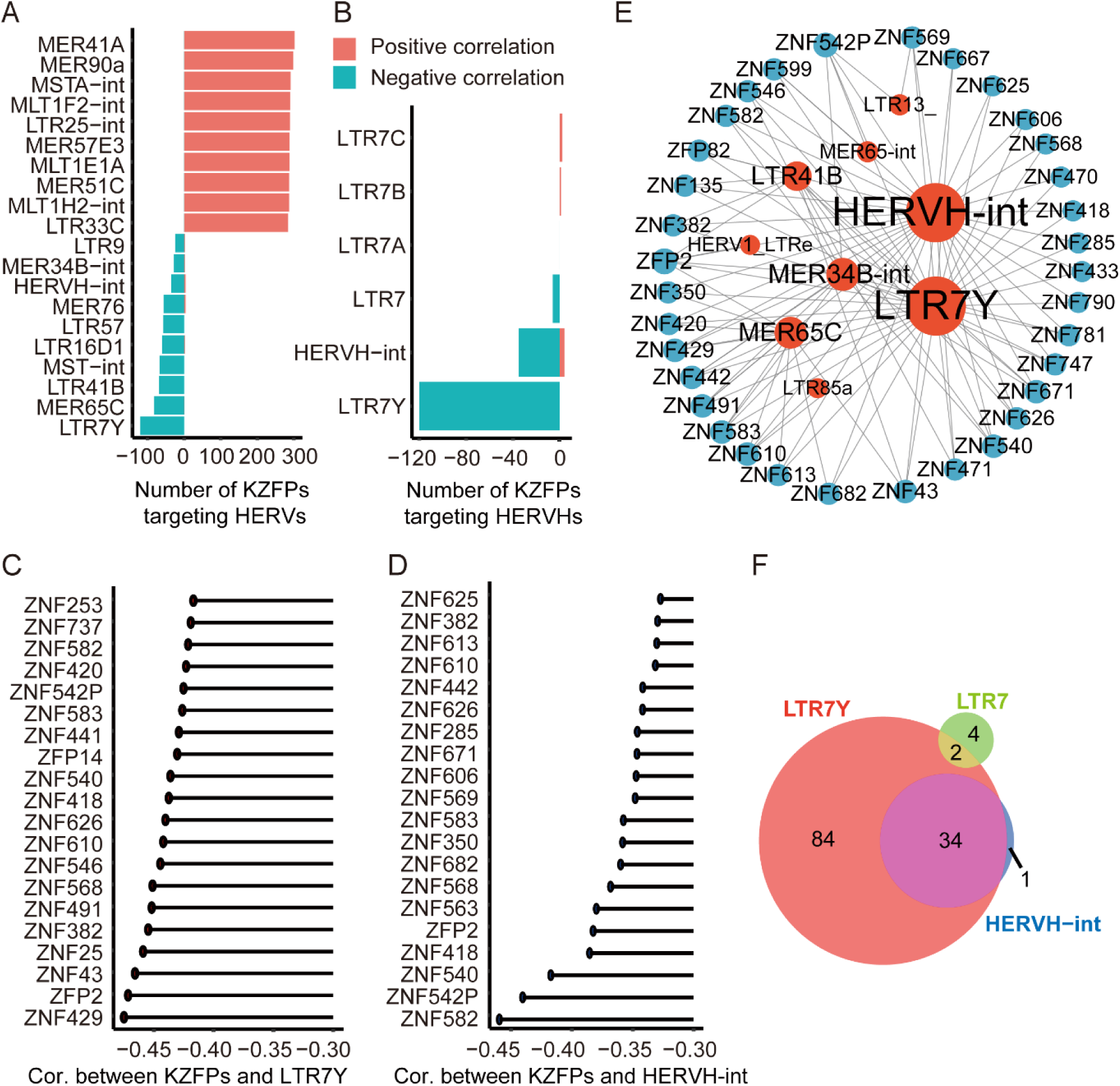
Case study: A pan-cancer survey of the correlations between KZFPs’ and HERVs’ expressions. (A) Bar plot showing the top 10 HERV subfamilies with the highest counts of positively or negatively correlated KZFPs, where negative or positive numbers represent the count of negatively or positively correlated KZFPs. (B) Bar plot showing the number of KZFPs positive or negatively correlated with the HERVH members. (C-D) The top 20 KZFPs negatively correlated with LTR7Y and HERVH-int. Cor. is the abbreviation of correlation. (E) Integrated regulatory network of KZFPs targeting HERVs, where blue dots represent KZFPs, red dots represent HERVs, and dot size indicates the number of connections. (F) Venn diagram displays the intersection of KZFPs negatively correlated with the HERVH members.

## Discussion

One unmet need of scientists from the HERVs research field is to easily screen abnormally activated HERVs in cancer samples to identify those involved in tumorigenesis therefore decode their functions. Exploiting a large collection of RNA-seq expression profiles of HERVs, ERVcancer helps to fill this gap in a straightforward manner. ERVcancer incorporates not only 7112 tissue samples from the TCGA project, but also 932 cancer cell samples from the CCLE project and 151 human early embryonic and embryonic stem cell samples from the GEO database. To our knowledge, this is the first online tool specifically designed to analyze HERVs’ expression in cancer cells and tissues. Through ERVcancer, users without any computational programming skills can easily explore large datasets, pose specific questions, and test hypotheses. For instance, users can readily identify that HERV-Fc2 subfamily is overexpressed in several cancers, such as breast invasive carcinoma, bladder urothelial carcinoma, head and neck squamous cell carcinoma, and liver hepatocellular carcinoma. Additionally, users can easily find HERVs that exhibit a significant association with patient survival, such as the HERVH-int in colorectal cancer and kidney renal clear cell carcinoma. Moreover, users can quickly identify HERVH-int is overexpressed in colon adenocarcinoma, with its most correlated coding gene being ESRG (Figure 3H). We believe this web recourse will accelerate the identification of HERVs that bear the potentials to be cancer biomarkers or even new therapeutic targets in future studies.

Previous reports have shown that KZFPs are widely recognized as transcriptional silencers against HERVs (Imbëault et al., 2017; Helleboid et al., 2019; Yang et al., 2022). For instance, ZFP961 specifically targets the Primer Binding Site (PBS)-Lys-containing ERVK for repression (Yang et al., 2022). We conducted a thorough investigation into the correlations between the expression profiles of KZFPs and HERVs in cancer. Surprisingly, we discovered that certain HERVs subfamilies were positively correlated with many KZFPs, while others showed predominant negative correlations. These findings suggest that the regulation of KZFPs on HERVs might be heterogeneous among different subfamilies. We then narrowed our focus to the HERVs that showed mostly negative correlations with KZFPs, which implies that they are often suppressed by these KZFPs. Notably, members of the HERVH family, which is tightly linked to the pluripotency network (Wang et al., 2014; Ai et al., 2022), were identified as “hub” HERVs that are targeted by KZFPs. Some of the HERVH-related KZFPs, such as ZNF382, is often down-regulated due to promoter methylation (Chen et al., 2020). It is compelling to speculate that down-regulation of ZNF382 may promote malignancy by derepressing the HERVH subfamily, reminiscent to our previous discovery which denotes that mutation of a tumor suppressor ARID1A reactivates a group of HERVH elements, enabling cancer cells to exploit and repurpose developmental programs to promote malignancy (Li et al., 2022; Yu et al., 2022). Therefore, the HERVH subfamily might serve as a general signal amplifiers to boost and strengthen the initial oncogenic signal during carcinogenesis (Li et al., 2022).

In summary, ERVcancer represents a straightforward and powerful web resource for investigating HERVs’ activation across many cancer types, complementing other available resources like ERVmap and repeatMasker. We hope that our platform can facilitate researches in this emerging field. However, the current version of the database has several limitations. Currently, a locus-specific atlas of HERVs’ expression profiles in cancer tissues and cells is unavailable, which limits our analysis only to the transcriptome data, leaving out all the locus-specific features such as local epigenetic landscapes including DNA methylation and histone modifications, as well as binding of key regulatory proteins.

Moving forward, we plan to continuously update our web resource with more comprehensive datasets. First, we will pinpoint all the HERVs loci that are unambiguously activated in different cancer types using only the uniquely mapped sequencing reads. On top of that, we will then include DNA methylation datasets from TCGA (https://portal.gdc.cancer.gov), reduced representation bisulfite sequencing data from CCLE (Ghandi et al., 2019), ChIP-seq data for KZFPs and other proteins from ENCODE and GEO (Dunham et al., 2012; Imbëault et al., 2017), as well as RNA-seq data for more cancer types and tissues. We envision that the next version of ERVcancer will allow users to query HERVs’ activation across major cancer types at the locus level, correlating HERVs’ activity with local epigenetic features and nearby gene expressions. Although it is still in its nascent stages of development, we believe that the exploration of HERVs in cancers holds significant potential in clinics.

## Materials and Methods

### Database and web interface

The ERVcancer database is implemented on CentOS Linux 7 using the Django web framework (https://www.djangoproject.com/), which is based on Python (V3.9.15). Datasets were organized into tables using SQLite3 (https://www.sqlite.org/). The web interface was developed using HTML, CSS, and JavaScript. Tools performing statistical tests were driven using Python library SciPy (https://scipy.org/) or using Matplotlib for graphics (https://matplotlib.org/).

### HERVs subfamily general information

We obtained the classification and annotation information of HERVs from UCSC (https://genome.ucsc.edu/cgi-bin/hgTables) (Lee et al., 2022), and sourced the consensus sequence (for GRCh38/hg38) of HERVs from RepBase (https://www.girinst.org/repbase/) and Dfam (https://www.dfam.org/) (Bao et al., 2015; Hubley et al., 2016).

### RNA-seq data analysis

For each RNA-seq data, firstly, FASTQ-formatted reads were trimmed adapter and cleaned using “trim_galore” (V0.6.0, http://www.bioinformatics.babraham.ac.uk/projects/trim_galore/) with default parameters. Then RNA-seq reads were aligned to the reference genome (GRCh38/hg38) using “STAR” (V2.5.3a) (Dobin et al., 2013); we included multiple-mapping reads, assigned them on random and reported only one alignment, with “-outMultimapperOrder Random -- outSAMmultNmax 1” option. Subsequently, RNA-seq reads mapped on HERVs subfamilies were counted using Subread “featureCounts” based on repeats annotation files downloaded from UCSC (Liao et al., 2014; Lee et al., 2022), with “--M” option. As for BAM-formatted RNA-seq data from TCGA which already reported all alignment for multiple-mapping reads using “STAR” (https://portal.gdc.cancer.gov/), we used “featureCounts” to assign fractional counts to repeat regions (Liao et al., 2014), with “--M --fraction” option, i.e. each reported alignment from a multi-mapping read will carry a fractional count of 1/x, instead of 1, where x is the total number of alignments reported for the same read.

### Differential expression analysis

For each HERV, differential expression analysis between tumor and normal tissues was performed based on the negative binomial distribution using the R package “DESeq2” (V1.22.2) (Love et al., 2014).

### Survival analysis

To assess the impact of HERVs on the prognosis of specific cancer, tumor samples were categorized to high and low-expression groups. Survival analysis between the high and low-expression groups was conducted using the “KaplanMeierFitter” function from the “lifelines” Python package (https://lifelines.readthedocs.io/).

### Correlation analysis

For each cancer type, pairwise correlations between the expression of HERVs and genes (or between HERVs) were calculated and visualized using the “regplot” function from “Seaborn” Python package (https://seaborn.pydata.org/).

### ZFP and HERVs correlation analysis

First, a list of KZFPs was obtained from de Tribolet-Hardy et al.(de Tribolet-Hardy et al., 2023). Second, using data from 6478 tumor tissue samples from TCGA, the standardized expression of 391 KZFPs and 580 HERVs subfamilies were merged. Third, the correlation coefficient and p value of each KZFP and HERV subfamily were calculated. When the absolute value of the correlation coefficient is larger than 0.3 (Ito et al., 2020), the corresponding KZFP and HERV subfamily pair was considered as statistically significant and recorded.

## Data availability

All processed data can be freely retrieved from the ERVcancer database for academic use. The major code or scripts are available in http://github.com/maosong327/ERVcancer.

## CRediT authorship contribution statement

Xiaoyun Lei: Methodology, Validation, Formal Analysis, Investigation, Data Curation, Writing-Original Draft, Visualization. Song Mao: Methodology, Validation, Software, Formal Analysis, Investigation, Data Curation, Writing-Original Draft, Visualization. Yinshuang Li: Methodology, Validation, Formal Analysis, Data Curation, Visualization. Shi Huang: Resources. Jinchen Li: Methodology, Resources. Wei Du: Resources. Chunmei Kuang: Writing-Original Draft, Supervision. Kai Yuan: Conceptualization, Resources, Writing-Review & Editing, Supervision, Project Administration, Funding Acquisition.

## Acknowledgements

We gratefully acknowledge Dr. Guihu Zhao and members of the Yuan Lab for inspiring discussions and technical assistance. This project has been supported by the National Natural Science Foundation of China (grants 32370821, 32170821, and 92153301 to K.Y), National Key Research and Development Program of China (2021YFC2701200), Department of Science & Technology of Hunan Province (grants 2023RC1028, and 2023SK2091 to K.Y), Middle/Young aged Teachers’ Research Ability Improvement Project of Guangxi Higher Education (grant 2024KY0111 to X.L), and Joint Project on Regional High-Incidence Diseases Research of Guangxi Natural Science Foundation (grant 2023JJB140356 to X.L).

---

**Figure S1.**
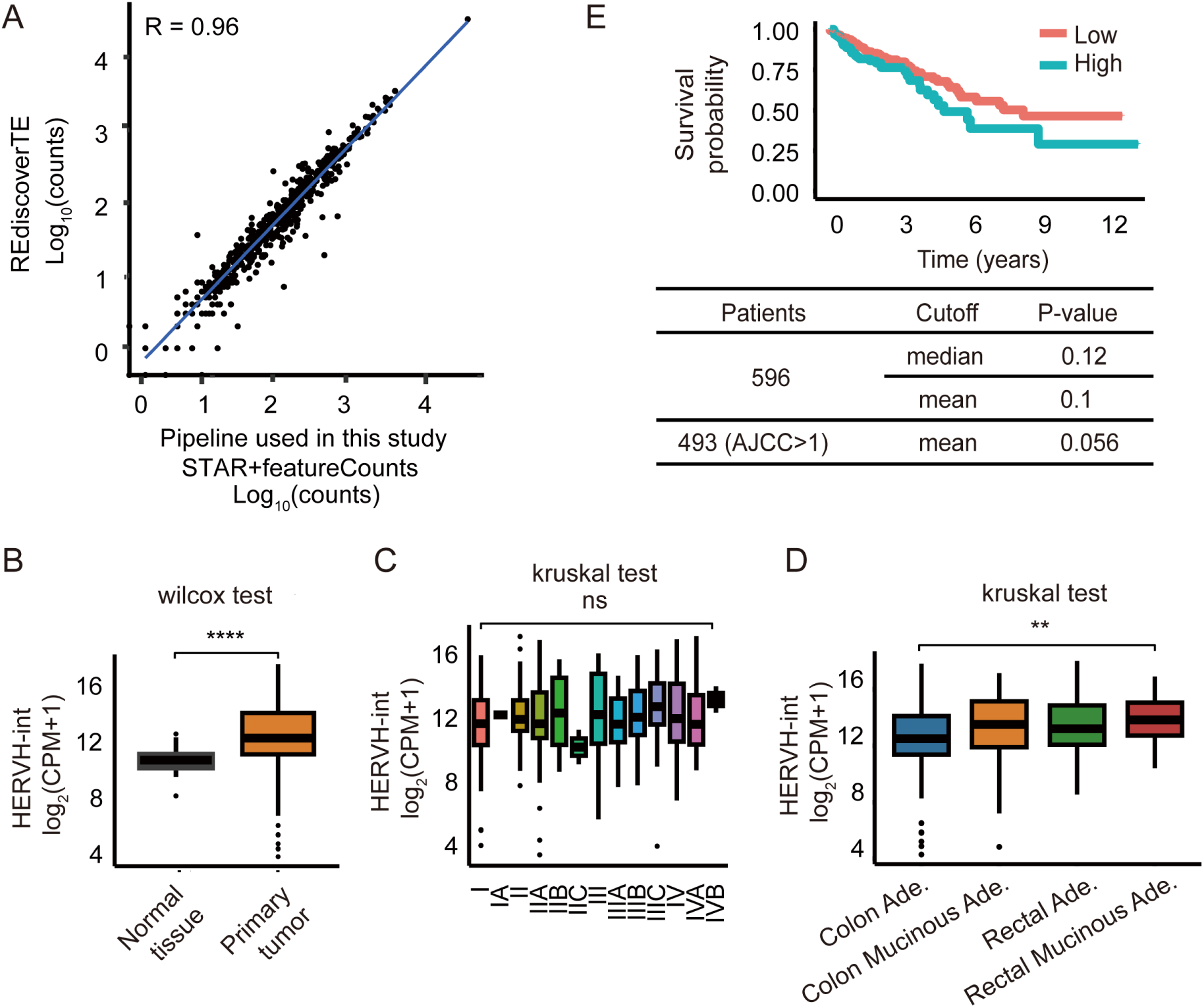
Benchmarking the accuracy of HERVs’ quantification and analysis by comparing our pipeline with REdiscoverTE. (A) Scatter plot illustrates the performance of the two pipelines in quantifying HERVs using a sample (TCGA-EI-7002-01A-11R-1928-07) from The Cancer Genome Atlas Colorectal Adenocarcinoma (TCGA-COREAD). Each dot represents an HERV subfamily and the value corresponding to the quantified counts. Performance accuracy is measured in terms of Pearson correlation coefficient R. (B-E) The same analysis was performed using the quantification results from REdiscoverTE, with TCGA-COREAD (contains 51 solid normal tumor tissues and 631 primary tumor tissues) as an example, corresponding to the results generated using our pipeline in Figure 3C-3F. Differential expression analysis of HERVH-int was performed among normal and cancer tissues (B), different tumor stages (C), or histological types (D). Overall survival analysis was performed according to the HERVH-int expression levels (E).

